# Psilocybin rapidly, but not immediately, reverses reward learning deficits in a durable manner in an inflammatory rat model of depressive symptoms

**DOI:** 10.64898/2026.01.14.699553

**Authors:** Justyna K Hinchcliffe, Christopher W Thomas, Gary Gilmour, Emma SJ Robinson

## Abstract

The serotonergic psychedelic, psilocybin, shows potential for rapid and sustained antidepressant effects but the underlying mechanisms remain unknown. Using a chronic interferon-alpha–induced rat model of depression, we show acute psilocybin (0.3 mg/kg) reverses impaired reward-induced biases within 24hrs, with effects enduring for at least 7 days. This suggests psilocybin can restore blunted reward processing, an effect which could significantly contribute to its sustained antidepressant effects.

## Main text

Recent clinical trials with serotonergic psychedelics suggest these may also induce antidepressant effects within 24hrs of a single administration, with effects potentially lasting for several months in some patients^1-3^. The mechanisms underlying these effects are poorly understood.

It is not possible to directly relate behavioural readouts in animal models to clinical rating scales used as regulatory endpoints in clinical trials. However, some rodent behavioural tasks can quantify cognitive and affective domains of impairment relevant to MDD^4,5^. These tasks exhibit good properties of construct validity in comparison to their human homologues. An example is the probabilistic reward learning task where studies in MDD patients have found blunted reward learning^6^. Reverse translation of these tasks for rodents led to establishment of a foraging-based reward learning assay (RLA)^7^. Studies using the RLA in rodent disease models have found impaired reward-induced biases similar to blunted reward learning observed in depressed patients^7-10^.

The current study used a chronic interferon-alpha (IFNA) model of inflammation-induced depression to generate a phenotypic model to test the acute and sustained effects of psilocybin on reward learning^7^. IFNA was chosen as a model with substantial construct validity, with evidence that 30% - 50% of patients treated with IFNA for hepatitis C develop symptoms of MDD^11^. Animals (3 cohorts of 16 male Lister-hooded rats, Envigo, ∼300g at start of training) were first trained and tested in the RLA (see Figure 1A and 1B, online methods), to establish a baseline reward-induced bias, then split into three groups for chronic treatment (for timeline see Figure 1C). Only male rats were used for practical reasons however previous studies have not found any signficant sex differences in the RLA^12^. Each RLA test was carried out using the same protocol (see online methods). Animals were presented with two ceramic bowls containing different digging media, one of which concealed a food reward. To generate specific cue-reward memories, rewarded digging substrates, either A or B, were used as the cues to predict either a high or low value reward, with a third substrate C providing a blank, unrewarded substrate. Each animal underwent independent learning sessions where they were presented with A vs C or B vs C on different days. A total of 4 learning sessions were used with 2 x A vs C and 2 x B vs C and A and B paired with the high or low reward, counter-balanced across subjects. To quantify reward-induced bias, a choice test was used on day 5/6 where substrates A and B were presented together for the first time over 30 randomly reinforced trials. All cohorts exhibited a reward-induced positive bias following training making more choices for the high reward-paired substrate (p<0.001, one-sample t-test, Supplementary Figure S1). Animals were then split into three groups with two groups treated once daily for 14 days with interferon alpha (100 units/kg, IP, 4-5pm) to induce a depression-like phenotype and one group treated with vehicle. Treatments were continued throughout testing. To investigate the effects of psilocybin (COMP360, 0.3mg/kg, IP) on the IFNA-induced impairment in reward learning, animals were treated with either psilocybin or vehicle and tested in the RLA 1hr and 24hrs post administration. A new RLA was then performed the following week with the pairing sessions and choice test run post-psilocybin treatment (see Figure 1C for timeline). Data were analysed using a one- or two-way ANOVA with post-hoc pairwise comparisons were made following a main effect. % Reward-induced bias was analysed using a one sample t-test against a theoretical mean of 0% bias.

**Figure 1.**
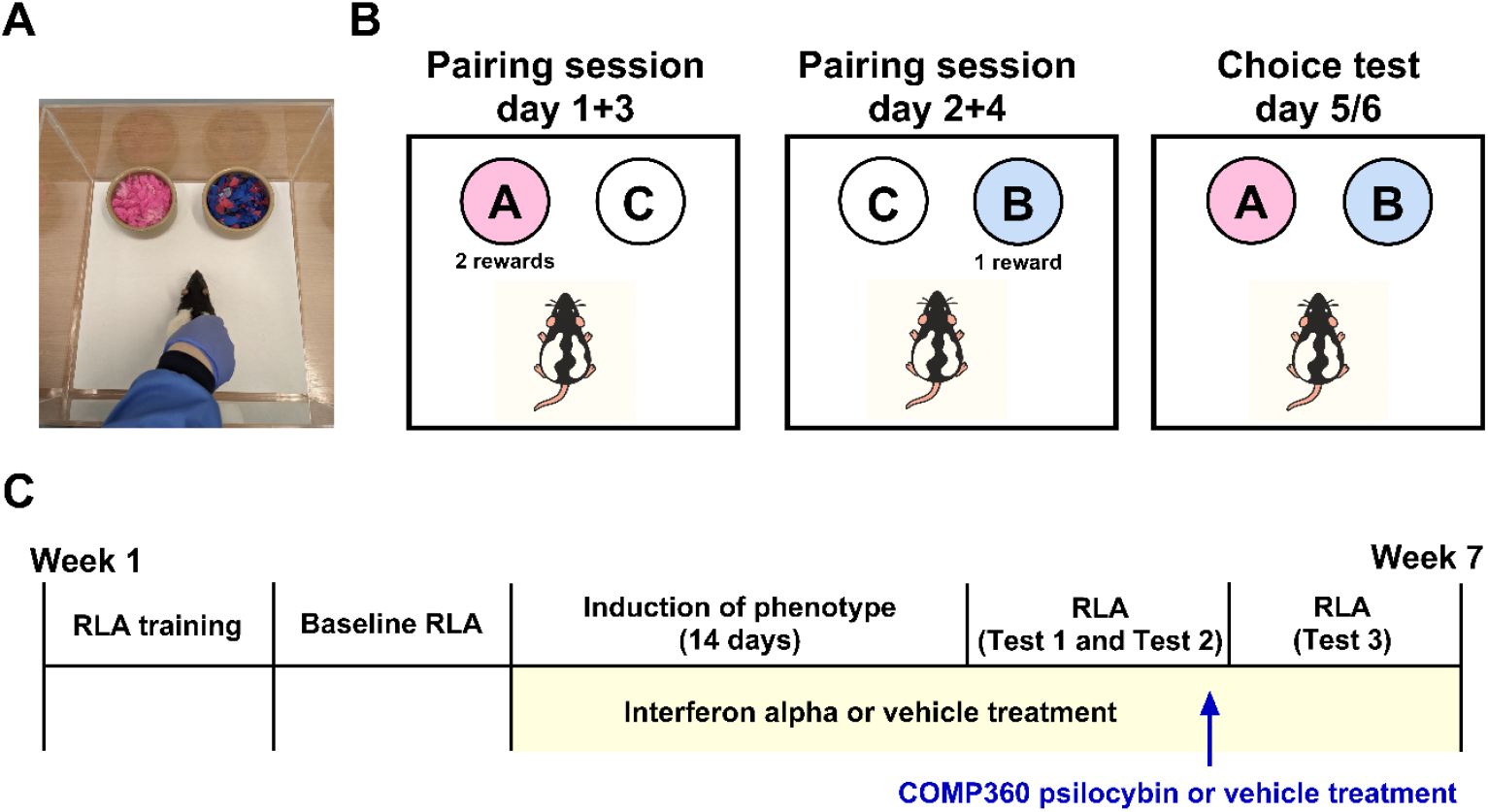
Reward learning assay. **A**. Photograph illustrating the RLA arena with bowls and rat position. **B**. Overview of the reward learning assay protocol. **C**. Experimental timeline.

Relative to the IFNA-vehicle group, psilocybin treated animals showed a reversal of the reward learning impairment at 24hrs and normalised reward learning for up to 7 days. For the study involving the repeated choice test at 1hr and 24hrs, there was a significant time × treatment interaction (F2,45=4.09, p=0.0234, Fig. 2A), indicating that treatment effects differed across time. There was a main effect of treatment (F2, 45=19.55, p<0.0001, N=16, Fig. 2A) but no main effect of time (F1,45=0.97, p=0.3304, Fig. 2A). Post hoc pairwise comparisons using Sidak’s correction showed that all animals treated with IFNA and acute vehicle prior to testing exhibited the predicted impairment in reward learning at both 1hr (p<0.0001) and 24hrs (p=0.0012). In the psilocybin treated group at 24hrs, the reward learning deficit was significantly attenuated (p=0.0170) with animals exhibiting a significant reward-induced positive bias (one sample t-test, t15=4.948, p=0.0002). The IFNA-psilocybin group were not significantly different (p=0.3243) from the vehicle-vehicle group, indicating a full amelioration of the IFNA-induced reward deficit (Fig 2A). When animals were tested in a new RLA (Test 3) with the choice test carried out 7 days following psilocybin administration, there was a main effect of treatment (One-way ANOVA, F2,29=4.188, p=0.0252, N=10-12, Fig. 2B) with both the vehicle-vehicle groups and interferon-alpha-psilocybin groups developing a reward-induced positive bias (one sample t-test, p=0.0138 and p=0.0002, respectively) and a difference between the IFNA-vehicle group and IFNA-psilocybin group meaning the effects on reward learning were sustained for at least 7 days (p=0.0291). The impaired reward-induce bias, relative to vehicle-vehicle control, was observed for the IFNA-vehicle group in all three of the post-treatment choice tests. No other effects of IFNA or psilocybin treatment during the pairing sessions or choice test were observed (Supplementary Tables S3, S4).

**Figure 2.**
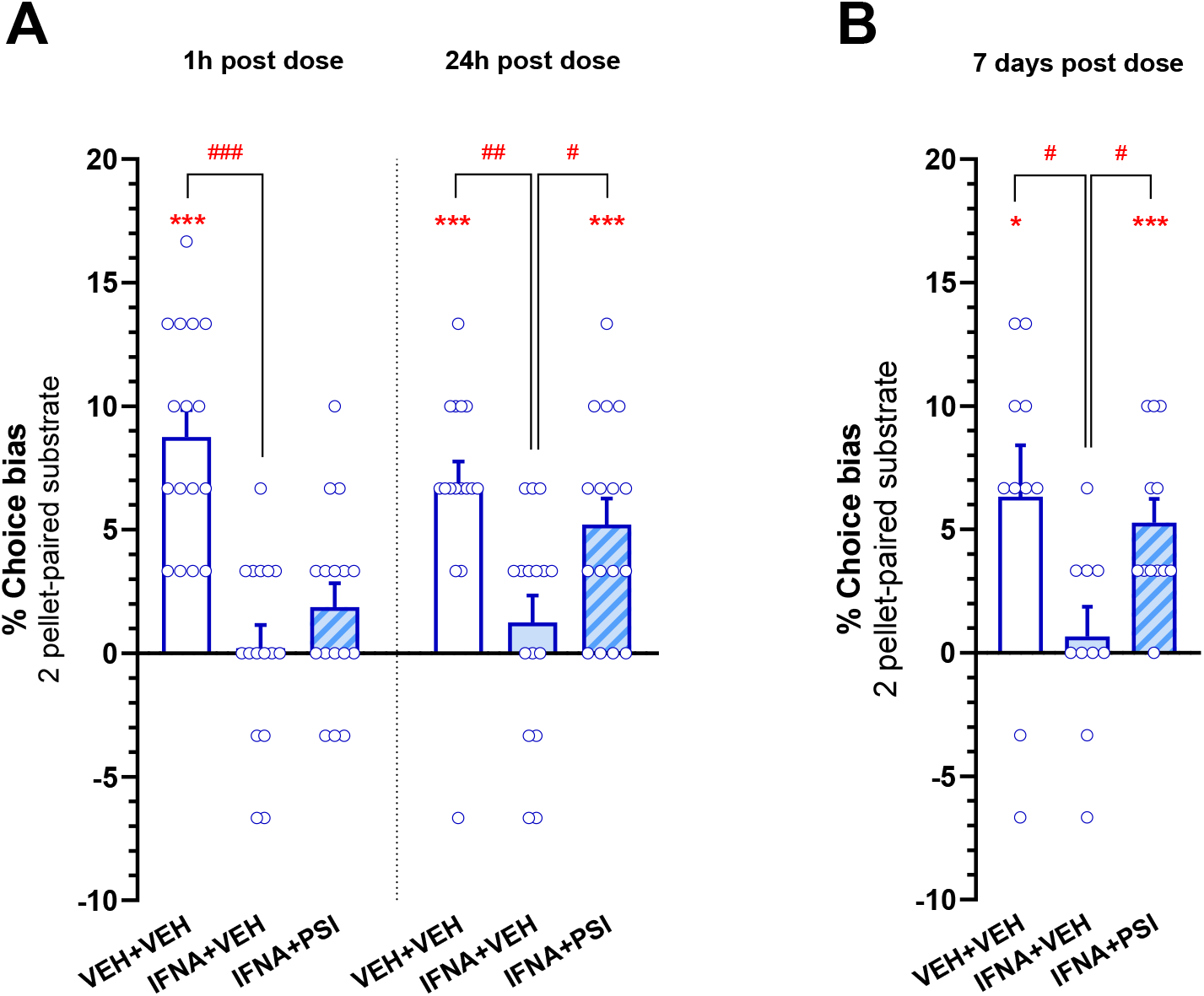
Acute psilocybin induces rapid and sustained amelioration of reward learning deficits. There was no effect of psilocybin at 1hr post-treatment but the blunted reward learning induced by IFNA was attenuated at 24hrs (panel A) post-treatment. The effects of psilocybin were sustained for at least 7 days with a similar attenuation of the IFNA-induced impairments in reward learning observed when a new RLA was performed with the choice test 7 days following treatment with psilocybin or vehicle (panel B). Data shown as mean ± SEM and individual data points, N = 10-16 per group, *p<0.05, ***p<0.001 (one-sample t test versus theoretical mean of 0% choice bias), ^#^p<0.05, ^##^p<0.01, ^###^p<0.001 (post-hoc Sidak’s test). VEH+VEH: vehicle-vehicle group, IFNA+VEH: interferon alpha-vehicle group, INFA-PSI: interferon alpha-psilocybin group.

Impairments in reward processing are an important behavioural feature of MDD which can be quantified objectively in a similar manner in both humans and rodents, offering a translational approach to the study of antidepressant effects in this domain. Using a chronic IFNA inflammatory model to induce relevant reward learning impairments and a reversal learning assay to measure reward processing, we show that an acute dose of psilocybin can reverse impaired reward learning within 24hrs and sustain these effects for at least 7 days. Such sustained effects could be speculated to be indicative of potentially plastic responses in reward learning circuits. It interesting to observe that the effects of psilocybin were not immediately detectable in this study, i.e. within 1 h of dosing. This is an important practical point to consider for future work, as it is not always commonplace to test parameters of interest beyond the acute dosing period if no effects are immediately observed.

fMRI imaging studies in MDD patients performing neuropsychological tasks suggest impairments in reward learning are associated with blunted responses in the nucleus accumbens (NAc). These may involve NAc specific changes or result from maladaptive alterations in wider reward circuits and connections with regions such as the prefrontal cortex and amygdala where hyperactivity in MDD has been observed^13^. Impairments in reward learning are likely to underpin loss of interest in previously rewarding activities, a core DSM-V symptom of depression which is often resistant to treatment with conventional SSRIs^14^. The ability of ketamine to modulate this core domain in MDD is hypothesised to underlie its RAAD effects and efficacy in treatment resistant populations^15^. These current findings reveal that psilocybin can also rapidly reverse reward learning impairments and enduring changes at a behavioural level suggesting induction of long-term adaption in relevant neural circuits.

The ability of psilocybin to induce long term changes in symptoms of MDD after a single administration does not align with drug-receptor models of treatment efficacy where clinical effects are considered to be related to ongoing drug exposure and continued engagement of the receptor or target of interest. Using a translational rodent model, we have been able to show that an important behavioural domain in MDD is directly modulated by an acute dose of psilocybin. We have previously shown in the affective bias test that psilocybin can facilitate the re-learning of negatively biased, past experiences with a more positive affective valence involving the prefrontal cortex^16^. A role for neuroplasticity and changes in connectivity in neural networks in emotional circuits has been proposed based on both human imaging and animal studies^16-18^ but what has not been understood is how these relate to the symptoms of MDD. Further studies integrating this behavioural approach with circuit-based analyses are needed but these findings suggest acute psilocybin treatment can induce rapid and sustained changes in reward processing mechanisms relevant to blunted reward learning and the symptom of anhedonia in MDD.

## Acknowledgements

Funding for this research was provided by a BBSRC Industrial Partnership grant awarded to ESJR and GG (BB/V015028/1) and included financial and in kind contributions from Compass Pathways Ltd.

## Methods

### Animals and Housing

Three cohorts of male Lister Hooded rats (n = 16 per cohort; Envigo, UK, N=4-6 per treatment group in each replicate) were used (see Suppl. Table S1). One cohort is not included in the 7 day experiment due to an error. All rats weighed between 300-400g (age: 10-13 weeks) at the beginning of the training. This study only used male rats for practical reasons although previous studies suggest similar reward learning biases are observed in male and female animals^12^. Rats were pair-housed in enriched cages, under a 12:12 h reverse light–dark cycle (lights off at 08:00 h) and maintained at 21±1°C. Food was restricted to ∼90% of free-feeding weight provided after testing and water was available *ad libitum*. All procedures followed the UK Animals (Scientific Procedures) Act 1986 and were approved by the University of Bristol AWERB and the UK Home Office (PPL number P9B6A09A1).

### Reward Learning Assay (RLA)

The reward learning assay used a protocol with 4 pairing sessions and a choice test where animals remain in the same affective state throughout the one-week protocol and learnt to associate the one reward-paired digging substrate with a high (2 pellet) and the other with a low (1 pellet) reward, followed by a choice test (day 5/6), see Fig.1A. For the details of habituation, training and study design see Suppl. Materials and Fig.1B. Once animals were trained and baseline RLA confirmed correct task performance in each cohort, with a reward-induced positive bias observed at population level. Animals were then treated for two weeks with daily interferon-alpha or vehicle injections and tested in the RLA (Test 1) with new substrate-reward associations. To test acute effects on memory, on day 5 of RLA all animals 1h prior to the choice test were injected either with vehicle or COMP360 psilocybin. Animals were then re-tested in the same choice test (with the same substrate-reward associations) 24hrs later (Test 2) to investigate the sustained effects. During the 5 days of RLA protocol animals were continuously treated with IFNA or vehicle. Starting the following Monday (3 days post-psilocybin) animals underwent a new set of substrate-reward pairings (see Table S2) using the same RLA protocol (Test 3) with reward-induced positive bias tested 7 days post-psilocybin. For this test, only 10-12 animals were included in the analysis due to an experimenter error meaning one cohort were run using incorrect substrates. The study used a between subject design with animals randomly allocated to the following groups: group 1 - chronic vehicle plus acute vehicle (control), group 2 - chronic interferon-alpha plus acute vehicle (phenotype control), group 3 - chronic interferon-alpha plus acute psilocybin.

### Drugs

The drug used to induce reward learning deficits was interferon alpha (100units per kg, administered intraperitoneally, daily for 40 days between 3-5pm and psilocybin (COMP360, an investigational medicinal drug/product that does not have marketing authorization and is not approved for therapeutic use other than in a clinical trial environment) (0.3mg/kg, single IP injection) was used to test its effects of on reward learning. The doses for interferon alpha and COMP360 psilocybin were based on our previous studies^7,16^. In both experiments, a between-subject design fully counterbalanced experimental design was used, with the experimenters’ blind to all treatments and the data only decoded once all replicates had been completed.

### Data analysis

Data were analysed using IBM SPSS Statistics 31 (IBM, USA) and figures were created using GraphPad Prism 10.4.0 (GraphPad Software, USA). Choice bias score was calculated as the number of choices made for the 2 pellets-paired substrate divided by the total number of trials multiplied by 100 to give a percentage value. A value of 50 was then subtracted to give a score where a choice bias towards the 2 pellets-paired substrate gave a positive value and a bias towards the 1 pellet-paired substrate gave a negative value. Choice bias scores for Test 1 and Test 2 were analysed with a two-way mixed ANOVA with time as within-subjects factor and treatment groups as between-subjects factor. Choice bias scores for Test 3 (new substrate-reward associations) and response latency during the choice test were analysed utilising a One-way ANOVA with treatment as the between-subject factor. *Post-hoc* analysis comparisons were made using Sidak’s test. Positive or negative affective biases were also analysed using a one-sample t-test against a null hypothesised mean of 0% choice bias.

## Supplementary materials

### Additional information for materials and methods

#### Detailed Methods

##### Animals and housing

Three cohorts of male Lister Hooded rats (Envigo, UK) were used in these experiments (n=16 each cohort, for details, see Supplementary Table S1). All rats weighed between 300-400g (age: 10-13 weeks) at the beginning of the training. The sample size was chosen based on our previous affective bias test studies and a meta-analysis which demonstrated large effect size for the reward-induced bias in Sprague Dawley and Lister Hooded rats ^1-2^. The meta-analysis suggests similar affective biases and effects size observed in both strains and sexes, therefore, to reduce the numbers of animals in these studies we used male only rats. All animals were pair-housed in standard enriched laboratory cages (55×35×21cm) with aspen woodchip bedding, paper bedding, cotton rope, wood block, cardboard tube (Ø 8 cm diameter) and red Perspex house (30×17×10cm), under a 12:12h reverse light–dark cycle (lights off at 08:00h) and in temperature-controlled conditions (21±1°C). Rats were food restricted to approximately 90% of their free feeding weights matched to the normal growth curve (∼18 g of laboratory chow (Purina, UK) per rat was placed in their cage food hopper and all rats were fed once at the end of the experimental day) and provided with *ad libitum* water. The behavioural procedures and testing were performed during the animals’ active phase between 09:00h and 17:00h. All animals were given daily health and welfare checks by the animal facility technicians and the researchers. All experimental procedures were conducted in accordance with the UK Animals (Scientific Procedures) Act 1986 and were approved by the University of Bristol Animal Welfare and Ethical Review Body and UK Home Office (PPL number P9B6A09A1).

#### Affective Bias Test (ABT)

##### General protocol

###### Training

The ABT testing was carried out in a Perspex® arena (40×40cm) with two ceramic bowls (Ø 10cm) and a trio of digging substrates (reward-paired substrates - ‘A’ or ‘B’ versus unrewarded substrate - ‘C’, matched for digging effort and counterbalanced across subjects; for details, see Supplementary Table S2). Prior to ABT training animals underwent two habituation sessions to the ABT arena (first without bowls, substrate or reward and second with empty bowls); rats were individually placed into the arena and allowed to explore for 10min. Further training consisted of three digging training sessions (20 trials per session) with a bowl filled with increasing amounts of digging substrate (sawdust) and a food reward (45mg purified rodent tablets, Test Diet, Sandown Scientific, UK). On the first day of digging training, each rat was placed in the arena and given 30s to approach and explore the empty bowl (without substrate) containing two pellets per trial. When the pellets were found and consumed, the trial was completed, and the rat was removed from the arena and the pellets were replenished in the bowl. During the next digging training session, each rat was given 30s to explore the bowl and start digging for a single pellet buried within 1 cm of sawdust. Following 20 trials in which the pellet was found and eaten, each rat was moved onto the final training session in which a single pellet was buried within 2 cm of sawdust. Once each animal was able to find a pellet within 30s on 10 consecutive trials (within a maximum 20 trials), the digging training was complete.

Following the training sessions, animals underwent a discrimination session allowing them to explore two bowls with two novel digging substrates (reward-paired substrate with single pellet versus unrewarded substrate). On each trial, the animal was individually placed in front of the two bowls. Once the animal made a choice by starting to dig in one bowl, the other bowl was removed by the experimenter. Choice of the reward-paired substrate was marked as a ‘correct’ trial, digging in the unrewarded substrate was classified as an ‘incorrect’ trial and if an animal failed to approach and explore the bowls within 30s, the trial was recorded as an ‘omission’. Trials were continued until the rat achieved six consecutive correct choices for the reward-paired substrate. The discrimination session allowed us to confirm that the animals could achieve our learning criterion of six consecutive correct trials in less than 20 trials. Once animals successfully reached criteria in the discrimination session, they were considered trained and progressed to testing in the reward learning assay.

###### Testing

Firstly, the baseline RLA was carried out in each cohort to confirm that the animals were correctly performing the task and at population level reward-induced positive bias was observed before animals progressed to studies involving chronic interferon alfa treatment. In the RLA, a similar protocol to the ABT is used ^1-3^ with each week of four pairing sessions (day 1-4, one per day) to generate two independent cue-specific memories, and a choice test (day 5/6) but animals remain in the same affective state throughout and learn to associate the two reward-paired digging substrates (substrate ‘A’ or ‘B’, counter-balanced across subjects) with either a low (1 - 45mg reward pellet) or high (2 pellets) reward (for details see Figure 1). During the pairing sessions, each trial involved presenting the rat with a choice between two bowls containing two different digging substrates, one of which was reward-paired (and manipulation) and contained a single 45mg reward pellet, and the other of which was unrewarded (substrate ‘C’). Substrate C, unrewarded, was kept the same for all four pairing sessions and a reward pellet was crushed into the bowl and mixed within the substrate, to prevent choices based on odour. One of substrates ‘A’ or ‘B’ was presented during pairing sessions on days 1 and 3, and the other was presented on days 2 and 4, with order counterbalanced across subjects. All factors (i.e. bowl location, substrates, pairing sessions) were fully counterbalanced. The number of trials to reach the criterion (six consecutive correct choices for the reward-paired substrate within each pairing session), and latency to dig were recorded for each animal.

Affective biases generated by this protocol were quantified during the choice test on day 5 or 6 of the protocol when the two previously rewarded substrates (‘A’ and ‘B’) were presented at the same time for 30 trials. In order to keep rats motivated to continue choosing without providing new associative information, a single 45mg food pellet was placed in either bowl using a random schedule with a probability of one in three, so that rats randomly received a reward (i.e. substrate ‘A’ contained a pellet on 10 of the 30 trials, and likewise for substrate ‘B’; on no trials were both bowls baited). Both bowls also had a pellet crushed and placed in the substrate to reduce the likelihood of the animal using odour to find the reward. The animals’ choices and latency to dig were recorded.

#### Drugs

The drug used to induce reward learning deficits was interferon alpha (100units per kg, administered intraperitoneally, daily for 40 days between 4-5pm and COMP360 psilocybin (0.3mg/kg, administered intraperitoneally, single injection at t=-60 min. and t=-24hrs) was used to test its effects of on reward learning. Rat recombinant IFN-α was procured from Sigma– Aldrich (UK). The concentration of the drug, once reconstituted with 1 ml sterile distilled water, was 1 × 10^5^ units/ml. This was aliquoted into stock vials and stored at −80 °C until use. On the injection day, the stock solution was used to prepare working 100 units per kg. COMP360 psilocybin was supplied by COMPASS Pathways plc. All drugs were dissolved in vehicle solution, 0.9% sterile saline and were freshly prepared every day and they were administered in a dose volume of 1.0 ml/kg. The doses for interferon alpha and COMP360 psilocybin were based on our previous studies ^1-3^. Intraperitoneal injection (IP) procedures were done by using a low-stress, non-restrained method. All animals were habituated to the holding position required for IP dosing for five days prior to the experiments. In both experiments, a between-subject design fully counterbalanced experimental design was used, with the experimenters blind to all treatments and the data only decoded once all replicates had been completed.

#### Data analysis

Data were analysed using IBM SPSS Statistics 31 (IBM, USA) and figures were created using GraphPad Prism 10.4.0 (GraphPad Software, USA). Remaining behavioural parameters, for each animal, mean trials to criterion and latency to dig during ABT pairing sessions were analysed using a paired t-test comparison between control (1 pellet) and manipulation (high reward - 2 pellets) for each week and latency to dig during choice test were analysed using an One-way ANOVA with treatment as the between-subject factor. Analysis of the choice latency and trials to criterion was made to determine the presence of any non-specific effects of treatment i.e. sedation (see Table S3 and S4). A Shapiro-Wilk test was used to determine a normal distribution for the % Choice bias, trials to criterion, and mean latency to dig during pairing sessions and choice test.

**Figure S1.**
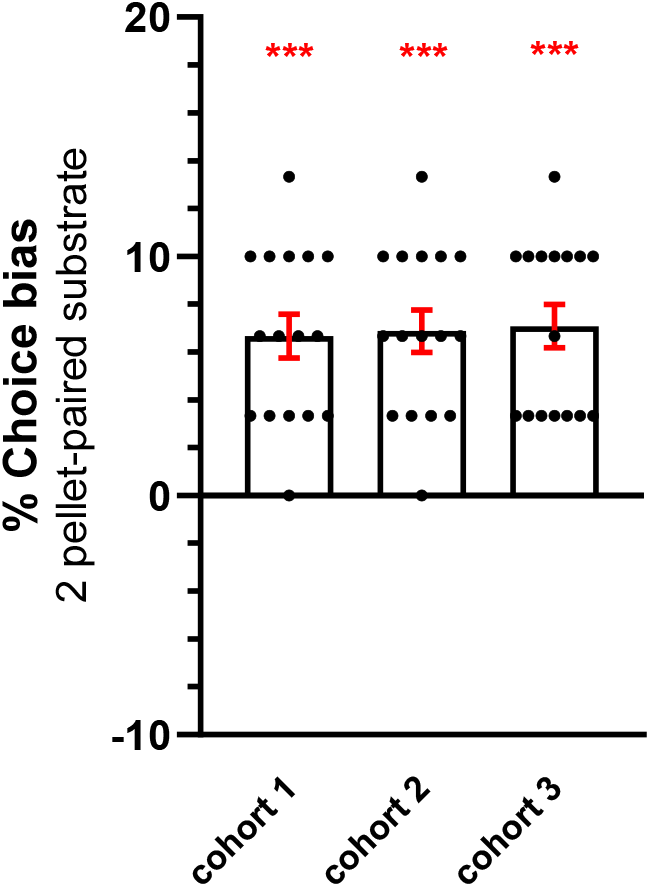
Control reward-learning assay data from each cohort used in the study before the start of treatment confirming successful learning. Data shown as mean ± SEM and individual data points, n = 16 per cohort, ***p<0.001 (one-sample t-test against theoretical mean of 0% choice bias).

**Table S1:** Summary of drug treatments in all cohorts.

| Cohort | Treatment | Dose | Route of administration | Treatment times |
| --- | --- | --- | --- | --- |
| 1, 2, 3, | Interferon alpha | 100units/kg | IP | Chronic daily (26 days) |
| 1,2, 3 | Psilocybin (COMP360) | 0.3mg/kg | IP | 60min., 24hrs and 7 days prior to choice test 1, 2 and 3 respectively |

**Table S3:** Pairing sessions data: number of trials to criterion and latency to dig. Data shown as mean (n=16 ^a^ and n=10-12 ^b^ animals/group) ± SEM averaged from the two pairing sessions for each substrate-reward association (1 pellet or 2 pellets). There were no significant effects during pairing sessions, either on response latency to dig or number of trials to criterion in any of the treatment groups.

| Treatment |  | Response latency |  | Trials to criterion |  |
| --- | --- | --- | --- | --- | --- |
|  |  | 1 pellet | 2 pellets | 1 pellet | 2 pellets |
| VEH-VEH | Week 1<sup>a</sup> | 2.6±0.1 | 2.7±0.2 | 6.3±0.1 | 6.4±0.1 |
| IFNA-VEH |  | 2.6±0.2 | 2.8±0.2 | 6.4±0.1 | 6.4±0.1 |
| IFNA-PSI |  | 2.8±0.2 | 2.6±0.2 | 6.4±0.1 | 6.3±0.1 |
| VEH-VEH | Week 2<sup>b</sup> | 2.4±0.2 | 2.2±0.1 | 6.4±0.1 | 6.4±0.1 |
| IFNA-VEH |  | 2.2±0.2 | 2.3±0.2 | 6.3±0.1 | 6.4±0.1 |
| IFNA-PSI |  | 2.0±0.1 | 2.3±0.1 | 6.3±0.1 | 6.4±0.1 |

**Table S2:** List of the substrates used in the experiments in all cohorts.

|  | Substrate 'A' | Substrate 'B' | Substrate 'C' |
| --- | --- | --- | --- |
| **Baseline RLA** | felt | shredded | exfoliating gloves |
| **RLA Test 1** | fur | polyester | pompoms |
| **RLA Test 2** | fur | polyester | pompoms |
| **RLA Test 3** | cellulose sponge | corrugated paper | perlite |

**Table S4:** Choice bias data: response latency to make choice. Data shown as mean (n=16 ^a^ and n=10-12 ^b^ animals/group) ± SEM of an individual latencies during 30 trials of the choice test. No significant difference in latency to make choice was observed following any of the treatments: vehicle – vehicle (0U/kg-0mg/kg, VEH-VEH), interferon alpha – vehicle (100U/kg-0mg/kg, IFNA-VEH) and interferon alpha – psilocybin (0U/kg-0.3mg/kg, IFNA-PSI).

| Treatment |  | Response latency (s) |
| --- | --- | --- |
| VEH-VEH | Test 1 <sup>a</sup> | 1.9±0.1 |
| IFNA-VEH |  | 1.8±0.1 |
| IFNA-PSI |  | 1.9±0.1 |
| VEH-VEH | Test 2 <sup>a</sup> | 2.1±0.0 |
| IFNA-VEH |  | 2.0±0.1 |
| IFNA-PSI |  | 2.0±0.1 |
| VEH-VEH | Test 3 <sup>b</sup> | 1.9±0.1 |
| IFNA-VEH |  | 1.9±0.1 |
| IFNA-PSI |  | 1.9±0.1 |

